# Disease variants alter transcription factor levels and methylation of their binding sites

**DOI:** 10.1101/033084

**Authors:** Marc Jan Bonder, René Luijk, Daria V. Zhernakova, Matthijs Moed, Patrick Deelen, Martijn Vermaat, Maarten van Iterson, Freerk van Dijk, Michiel van Galen, Jan Bot, Roderick C. Slieker, P. Mila Jhamai, Michael Verbiest, H. Eka D. Suchiman, Marijn Verkerk, Ruud van der Breggen, Jeroen van Rooij, Nico Lakenberg, Wibowo Arindrarto, Szymon M. Kielbasa, Iris Jonkers, Peter van ’t Hof, Irene Nooren, Marian Beekman, Joris Deelen, Diana van Heemst, Alexandra Zhernakova, Ettje F. Tigchelaar, Morris A. Swertz, Albert Hofman, André G. Uitterlinden, René Pool, Jenny van Dongen, Jouke J. Hottenga, Coen D.A. Stehouwer, Carla J.H. van der Kallen, Casper G. Schalkwijk, Leonard H. van den Berg, Erik. W van Zwet, Hailiang Mei, Yang Li, Mathieu Lemire, Thomas J. Hudson, the BIOS Consortium, P. Eline Slagboom, Cisca Wijmenga, Jan H. Veldink, Marleen M.J. van Greevenbroek, Cornelia M. van Duijn, Dorret I. Boomsma, Aaron Isaacs, Rick Jansen, Joyce B.J. van Meurs, Peter A.C. ’t Hoen, Lude Franke, Bastiaan T. Heijmans

**Affiliations:** Department of Genetics, University of Groningen, University Medical Centre Groningen, Groningen, The Netherlands; Molecular Epidemiology Section, Department of Medical Statistics and Bioinformatics, Leiden University Medical Center, Leiden, The Netherlands; Genomics Coordination Center, University Medical Center Groningen, University of Groningen, Groningen, The Netherlands; Department of Human Genetics, Leiden University Medical Center, Leiden, The Netherlands; SURFsara, Amsterdam, The Netherlands; Department of Internal Medicine, ErasmusMC, Rotterdam, The Netherlands; Sequence Analysis Support Core, Leiden University Medical Center, Leiden, The Netherlands; Medical Statistics Section, Department of Medical Statistics and Bioinformatics, Leiden University Medical Center, Leiden, The Netherlands; Department of Gerontology and Geriatrics, Leiden University Medical Center, Leiden, The Netherlands; Department of Epidemiology, ErasmusMC, Rotterdam, The Netherlands; Department of Biological Psychology, VU University Amsterdam, Neuroscience Campus Amsterdam, Amsterdam, The Netherlands; Department of Internal Medicine and School for Cardiovascular Diseases (CARIM), Maastricht University Medical Center, Maastricht, The Netherlands; Department of Neurology, Brain Center Rudolf Magnus, University Medical Center Utrecht, Utrecht, The Netherlands; Ontario Institute for Cancer Research, Toronto, Ontario, Canada M5G 0A3; Department of Medical Biophysics, University of Toronto, Toronto, Ontario, Canada M5S 1A1; Department of Molecular Genetics, University of Toronto, Toronto, Ontario, Canada M5S 1A1; Genetic Epidemiology Unit, Department of Epidemiology, ErasmusMC, Rotterdam, The Netherlands; Department of Psychiatry, VU University Medical Center, Neuroscience Campus Amsterdam, Amsterdam, The Netherlands

**Author notes:** Shared first (Marc Jan Bonder, René Luijk). Shared second (Daria V. Zhernakova, Matthijs Moed, Patrick Deelen, Martijn Vermaat). Shared second last (Aaron Isaacs, Rick Jansen, Joyce B.J. van Meurs). Shared last (Peter A.C. ’t Hoen, Lude Franke, Bastiaan T. Heijmans). Corresponding authors: Lude Franke and Bastiaan T. Heijmans.

## Abstract

Most disease associated genetic risk factors are non-coding, making it challenging to design experiments to understand their functional consequences^1,2^. Identification of expression quantitative trait loci (eQTLs) has been a powerful approach to infer downstream effects of disease variants but the large majority remains unexplained.^3,4^. The analysis of DNA methylation, a key component of the epigenome^5^, offers highly complementary data on the regulatory potential of genomic regions^6,7^. However, a large-scale, combined analysis of methylome and transcriptome data to infer downstream effects of disease variants is lacking. Here, we show that disease variants have wide-spread effects on DNA methylation *in trans* that likely reflect the downstream effects on binding sites of *cis*-regulated transcription factors. Using data on 3,841 Dutch samples, we detected 272,037 independent *cis*-meQTLs (FDR < 0.05) and identified 1,907 trait-associated SNPs that affect methylation levels of 10,141 different CpG sites *in trans* (FDR < 0.05), an eight-fold increase in the number of downstream effects that was known from *trans*-eQTL studies^3,8,9^. *Trans*-meQTL CpG sites are enriched for active regulatory regions, being correlated with gene expression and overlap with Hi-C determined interchromosomal contacts^10,11^. We detected many *trans*-meQTL SNPs that affect expression levels of nearby transcription factors (including *NFKB1, CTCF* and *NKX2–3*), while the corresponding *trans*-meQTL CpG sites frequently coincide with its respective binding site. *Trans*-meQTL mapping therefore provides a strategy for identifying and better understanding downstream functional effects of many disease-associated variants.

---

To systematically study the role of DNA methylation in explaining downstream effects of genetic variation, we analysed genome-wide genotype and DNA methylation in whole blood from 3,841 samples from five Dutch biobanks^12–16^ (Figure 1 and Extended Data Table 1). We found *cis*-meQTL effects for 34.4% of all 405,709 tested CpGs (n=139,566 at a CpG-level FDR of 5%, *P* ≤ 1.38 × 10^−4^), typically with a short physical distance between the SNP and CpG (median distance 10 kb, Extended Data Fig. 1). By regressing out primary meQTLs effect for each of these CpGs and repeating the *cis*-meQTL mapping, we observed up to 16 independent *cis*-meQTLs for these CpGs (Extended Data Table 2). In total, we identified 272,037 independent *cis*-meQTL effects. Few factors determine whether a CpG site shows a *cis*-meQTL effect except the variance in methylation level of the CpG site involved: for the top 10% most variable CpGs, 57.2% showed a *cis*-meQTL effect, dropping to only 8.1% for the 10% least-variable CpGs (Extended Data Fig. 2, Extended Data Fig. 3a). The proportion of methylation variance explained by SNPs, however, is typically small (Extended Data Fig. 3b). When accounting for this strong effect of CpG variation, we find only modest enrichments and depletions for *cis*-meQTL CpG sites when using CpG island (CGI) and genic annotation (Extended Data Fig. 3e) or when using annotations of biological function based on chromatin segmentations of 27 blood cell types (Figure 2a).

**Figure 1.**
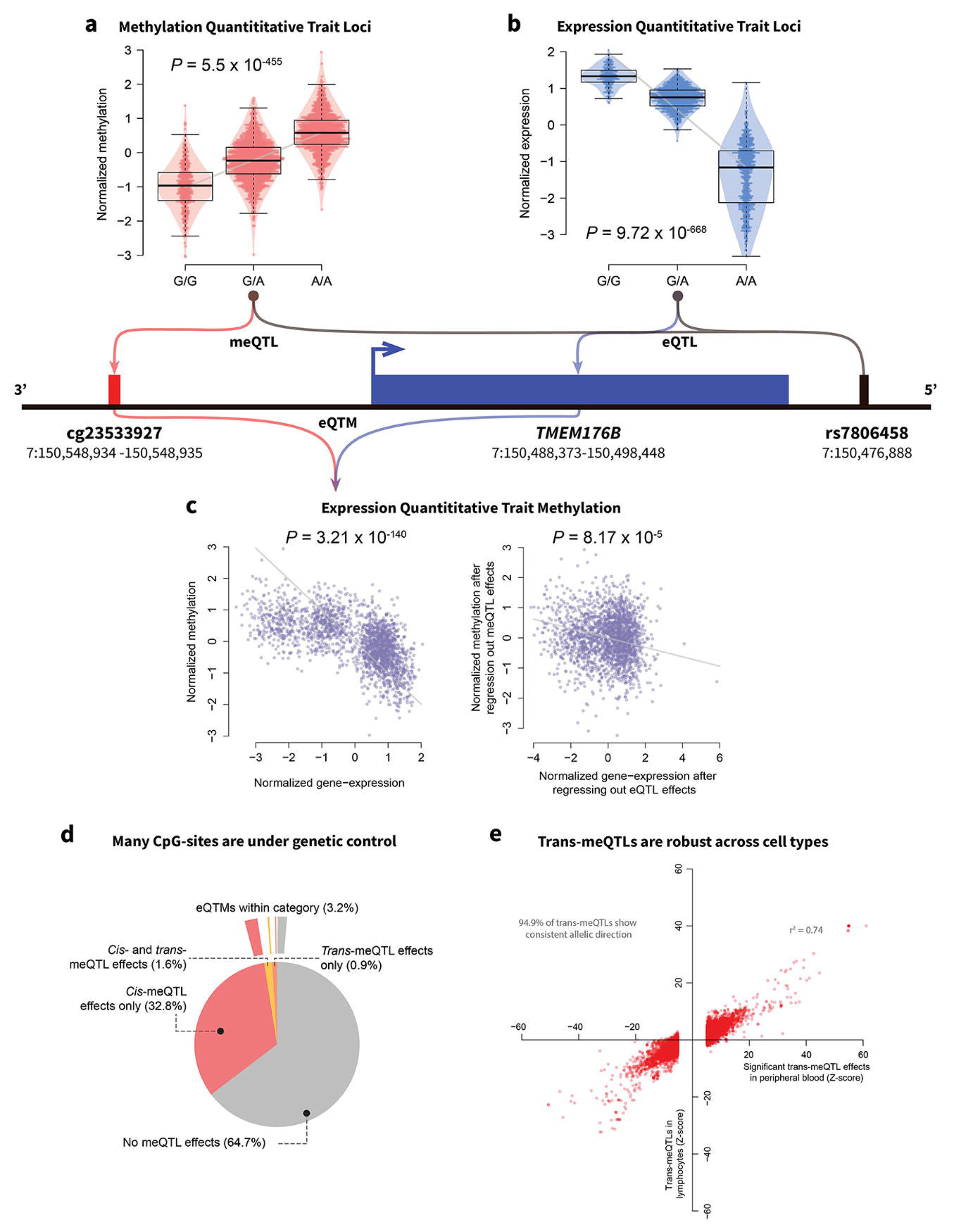
Overview of a genomic region around *TMEM176B*, where the relations between a SNP, DNA methylation at nearby CpGs, and the associations with the gene itself are shown. **a**, Illustration of a methylation Quantitative Trait Locus (meQTL) **b**, Illustration of an expression Quantitative Trait Locus (eQTL). **c**, Ilustration of methylation-expression association (eQTM). The figures show how correction for meQTLs may increase detection of such associations. The left plot shows the data before correction for *cis*-meQTLs, the corrected data in the right figure shows the meQTL-corrected methylation data. **d**, Two overlaid pie charts. The inner chart indicates the proportion of tested CpGs harboring meQTLs. Over 35% of all tested CpGs show evidence for harboring a meQTL, either *in cis* or *in trans*. The outer chart indicates what CpGs are associated with gene expression *in cis* (in total 3.2%). **e**, Replication of peripheral blood *trans*-meQTLs in lymphocytes.

**Figure 2.**
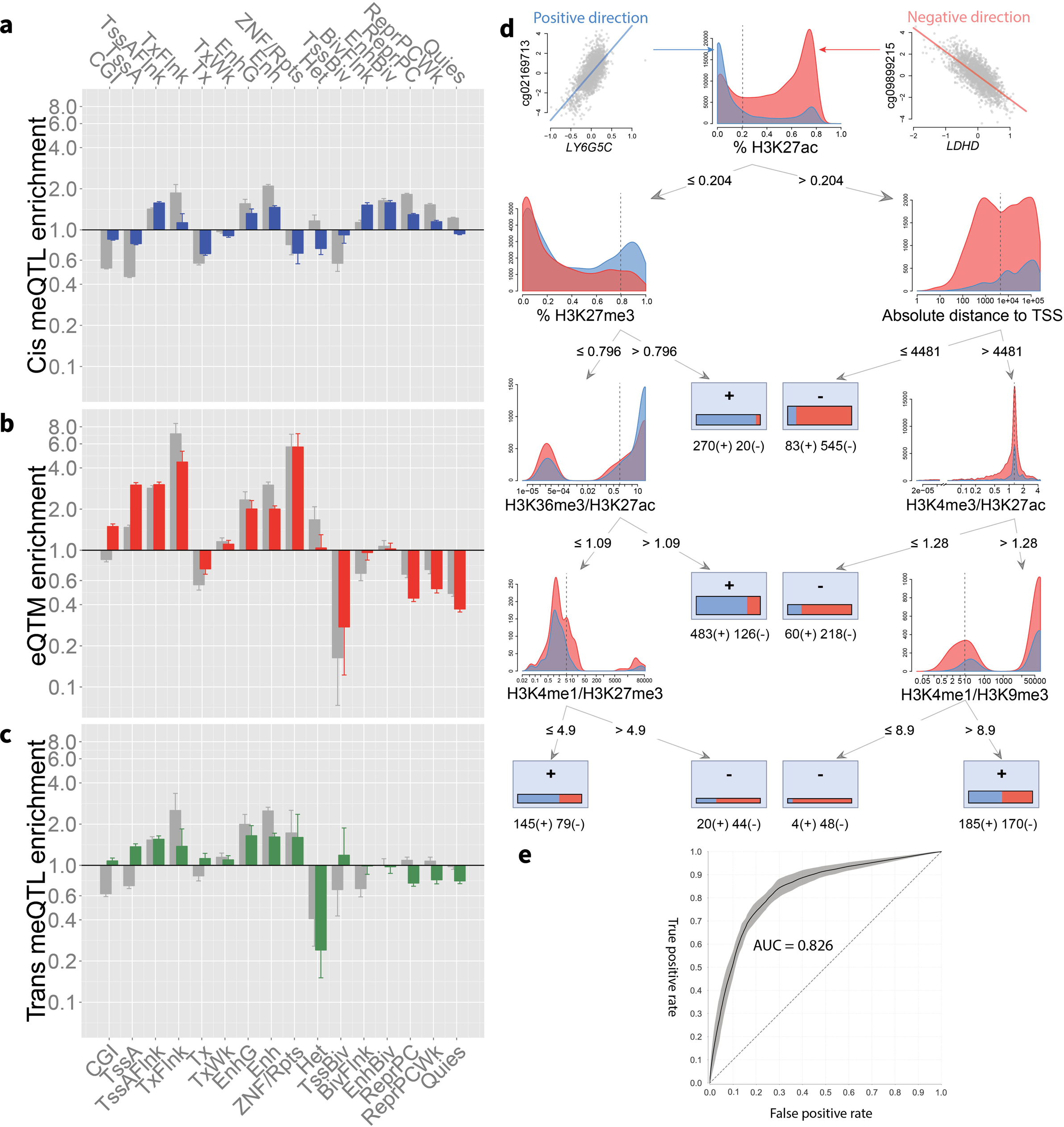
**a-c**, Over- or underrepresentation of CpGs for different predicted chromatin states for *cis*-meQTLs, *trans*-meQTLs and eQTMs. Grey bars reflect uncorrected enrichments, colored bars reflect enrichments after correction for factors influencing the likelihood of harboring a meQTL or eQTM, including methylation variability. Bar graphs show odds ratios and error bars (95% confidence interval). CGI: CpG island; TssA: Active TSS; TssAFlnk: Flanking active TSS; TxFlnk, Transcribed at gene 5’ and 3’; Tx: Strong transcription; TxWk: Weak transcription; EnhG: Genic enhancer; Enh: Enhancer; ZNF/Rpts: ZNF genes and repeats; Het: Heterochromatin; TssBiv: Bivalent/Poised TSS; BivFlnk: Flanking bivalent TSS/Enhancer; EnhBiv: Bivalent enhancer. **d**, Decision tree for predicting the effect direction of eQTMs. Each subplot shows the distributions for positive (blue) and negative (red) associations for that subset of the data. Dashed vertical lines indicate the optimal split used by the algorithm. The boxes in the leaves indicate the number of positive and negative effects in each of the leaves. **e**, Receiver operator characteristic curve showing the performance of the decision tree.

We contrasted these modest functional enrichments to CpGs whose methylation levels correlates with gene expression *in cis* (i.e. mapping expression quantitative trait methylations (eQTMs)), by generating RNA-seq data for 2,101 out of 3,841 individuals in our study. Using a conservative approach that maximally accounts for potential biases (i.e. *cis*-meQTL effects, *cis*-eQTL effects, batch effects and cell heterogeneity effects), we identified 12,809 unique CpGs that correlated to 3,842 unique genes *in cis* (CpG-level FDR < 0.05). eQTMs were enriched for mapping in active regions, e.g. in and around active TSSs (3-fold enrichment, *P* = 1.8 *×* 10^−91^) and enhancers (2-fold enrichment, *P* = 1.1 *trans*- 10^−139^, Figure 2b). Of note, the majority of eQTMs showed the canonical negative correlation with transcriptional activity (69.2%) but a substantial minority of correlations was positive (30.8%) in line with recent evidence that DNA methylation does not always negatively correlate with gene expression^17^. As expected, negatively correlated eQTMs were enriched in active regions like active TSSs (3.7-fold enrichment, *P* = 9.5 *×* 10^−202^). Positive correlations primarily occurred in repressed regions (e.g. Polycomb repressed, 3.4-fold enrichment, *P* = 5.8 *×* 10^−103^) (Extended Data Fig. 4). The sharp contrast between positively and negatively associated eQTMs, enabled us to build a model to predict the direction of the correlation. A decision tree trained on the strongest eQTMs (those with an FDR < 9.7*×*10^−6^, n=5,137) using data on histone marks and distance relative to gene, could predict the direction with an area under the curve of 0.83 (95% confidence interval, 0.78–0.87) (Figure 2d, e).

We next ascertained whether *trans*-meQTLs are biologically informative, since previous *trans*-eQTL mapping studies demonstrated that identifying *trans*-expression effects provide a powerful tool to uncover and understand downstream biological effects of disease-SNPs^3,8,9^. We focussed on 6,111 SNPs that were previously associated with complex traits and diseases (‘trait-associated SNPs’, see Methods and Extended Data Table 3). We observed that one-third of these trait-associated SNPs (1,907 SNPs, 31.2%) affect methylation *in trans* at 10,141 CpG sites, totalling 27,816 SNP-CpG combinations (FDR < 0.05, *P* < 2.6*×*10^−7^, Figure 3a),. This represents a 5-fold increase in the number of CpG sites affected as compared with a previous trans-meQTL mapping study^18^. We evaluated whether the GWAS SNP themselves were likely underlying the *trans*-effects or that the associations could be attributed to another SNP in moderate LD. Of the 1,907 GWAS SNPs with trans-effects, 1,538 (87.2%) were in strong LD with the top SNP (R^2^ > 0.8), indicating that the GWAS SNPs indeed are the driving force behind many of the *trans*-meQTLs. Of note, due to the sparse coverage of the Illumina 450k array, the true number of CpGs in the genome that are altered by these trait associated SNPs will be substantially higher. After the identification of the trans-meQTLs, we assessed if the *trans*-meQTLs also are present in expression. Out of the 2,889 testable trans-eQTLs we identified 8.4% of these effects, 91% of the cases the effect direction was consistent (Extended Data Table 4).

**Figure 3.**
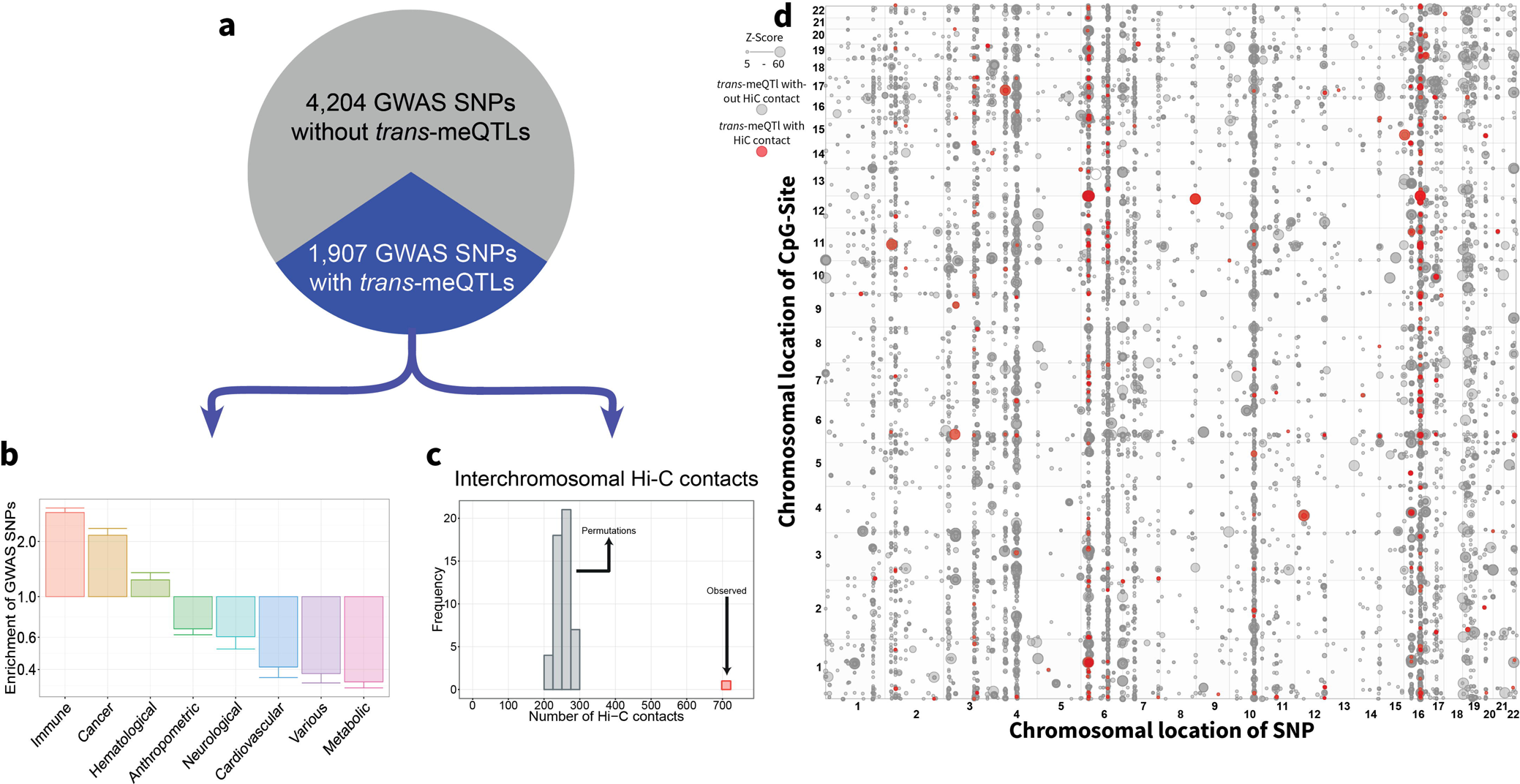
**a**, Distribution of tested trait-associated SNPs influencing DNA methylation *in trans*. Over 1,900 SNPs (31.2%) of all tested SNPs have downstream effects on DNA methylation. **b**, Overrepresentation of SNPs with *trans*-meQTLs in different GWAS trait categories, where the y-axis shows the odds ratio. **c**, Hi-C contacts are overrepresented among *trans*-meQTLs. Grey bars show the number of Hi-C contacts using permutated data, while the red bar reflects the actually observed number in our data. **d**, Dot-plot depicting the *trans*-meQTLs. The effect strength is reflected by the size of the dot. Red dots indicate an overlap with a Hi-C contact. Several SNPs with widespread *trans*-meQTLs show interchromosomal contacts genome-wide, further implicating an important role for those SNPs in the development of the associated trait.

To ascertain stability our *trans*-meQTLs, we performed a replication analysis in a the set of 1,748 lymphocyte samples^18^: of the 18,764 overlapping *trans*-meQTLs between the datasets that could be tested, 94.9% had a consistent allelic direction (Figure 1E). 12,098 *trans*-meQTLs were nominally significant (unadjusted *P* < 0.05), of which 99.87% had a consistent allelic direction. This indicates that the identified *trans*-meQTLs are robust and not caused by differences in cell-type composition. (Extended Data Table 5). To further ascertain the stability of the *trans*-meQTLs, we tested SNPs known to influence blood composition^19,20^ for effects on methylation *in trans*, finding these SNPs show no or only few *trans*-meQTLs whereas widespread trans-meQTL effects were to be expected if our analysis had not properly controlled for blood cell composition (Extended Data Table 6). Furthermore we linked our GWAS SNPs to the SNPs known to influence cell proportions and found that only 0.6% of the GWAS SNPs are in high LD with SNPs known to influence cell proportions. Lastly, we performed trans-meQTL mapping on uncorrected and cell type corrected data see supplemental results and Extended Data Table 7,8.

In contrast to *cis*-meQTL CpGs, *trans*-meQTLs CpGs show many functional enrichments: they are enriched around TSSs and depleted in heterochromatin (Figure 2c) and are strongly enriched for being an eQTM (1,913 CpGs (18.9%), 5.2-fold, *P* = 2.3 *00d7* 10^−101^). The 1,907 trait-associated SNPs that make up the *trans*-meQTLs were overrepresented for immune- and cancer-related traits (Figure 3b). The large majority of *trans*-meQTLs were inter-chromosomal (93%, 9,429 CpG-SNP pairs) and included 12 *trans*-meQTLs SNPs (yielding 3,616 unique CpG-SNP pairs) that each showed downstream *trans*-meQTL effects across all of the 22 autosomal chromosomes (i.e. *trans*-bands, Figure 3d).

We subsequently studied the nature of these *trans*-meQTLs. Using high-resolution Hi-C data^10^, we identified 720 SNP-CpG pairs (including 402 CpG sites and 172 SNPs) among the *trans*-meQTLs that overlapped with an inter-chromosomal contact, which is 2.9-fold more frequent than expected by chance (*P* = 3.7 *×* 10^−126^, Figure 3c, d). These Hi-C inter chromosomal enrichments were not confounded due to SNPs that gave *trans*-meQTLs on many CpG sites (i.e. *trans*-bands): when removing those *trans*-meQTLs from the analysis, Hi-C enrichments remained highly significant (*P* = 1.7*×*10^−61^). This indicates that some relationships between SNPs and CpGs *in trans* are explained by inter-chromosomal contacts. In order to characterize the 720 SNP-CpG pairs overlapping with inter-chromosomal contacts, we performed motif enrichments using three motif enrichment analyses (Homer, PWMEnrich, DEEPbind)^21–23^. These analyses identified that the 402 CpG sites frequently overlapped with CTCF, RAD21 and SMC3 binding sites (*P* = 2.3×10^−5^, *P* = 3.5*×*10^−5^ and *P* = 5.1*×*10^−5^, respectively), factors known to affect chromatin architecture^24,25^. This finding was confirmed by incorporating ChIP-Seq data on CTCF binding (1.8-fold enrichment, P = 5.2×10−7).

We next tested whether the *trans*-meQTLs reflected the effect of differential transcription factor (TF) binding of TFs that map close to the SNPs since TF binding has been implicated in demethylation and loss of TF occupancy with remethylation^6,7^. This suggests that if a SNP allele increases TF els *in cis*, that *trans*-meQTL effects are likely detectable, and that the SNP allele likely decreases methylation of these CpG sites. Indeed, we observed that if a SNP affects multiple CpGs sites *in trans* (at least 10, n=305) that the assessed allele often consistently increased or decreased methylation *in trans*, in the same direction for, on average, 76% of CpGs per *trans*-meQTL SNP (expected 50%, *P*=10^−111^; Figure 4a). This skew in allelic effect direction was present for 59.7% of the 305 SNPs with at least 10 *trans*-meQTL effects increasing to 95.2% for 104 SNPs with at least 50 *trans*-meQTL effects (binomial test P < 0.05), suggesting that differential TF binding may explain a substantial fraction of *trans*-meQTLs.

**Figure 4.**
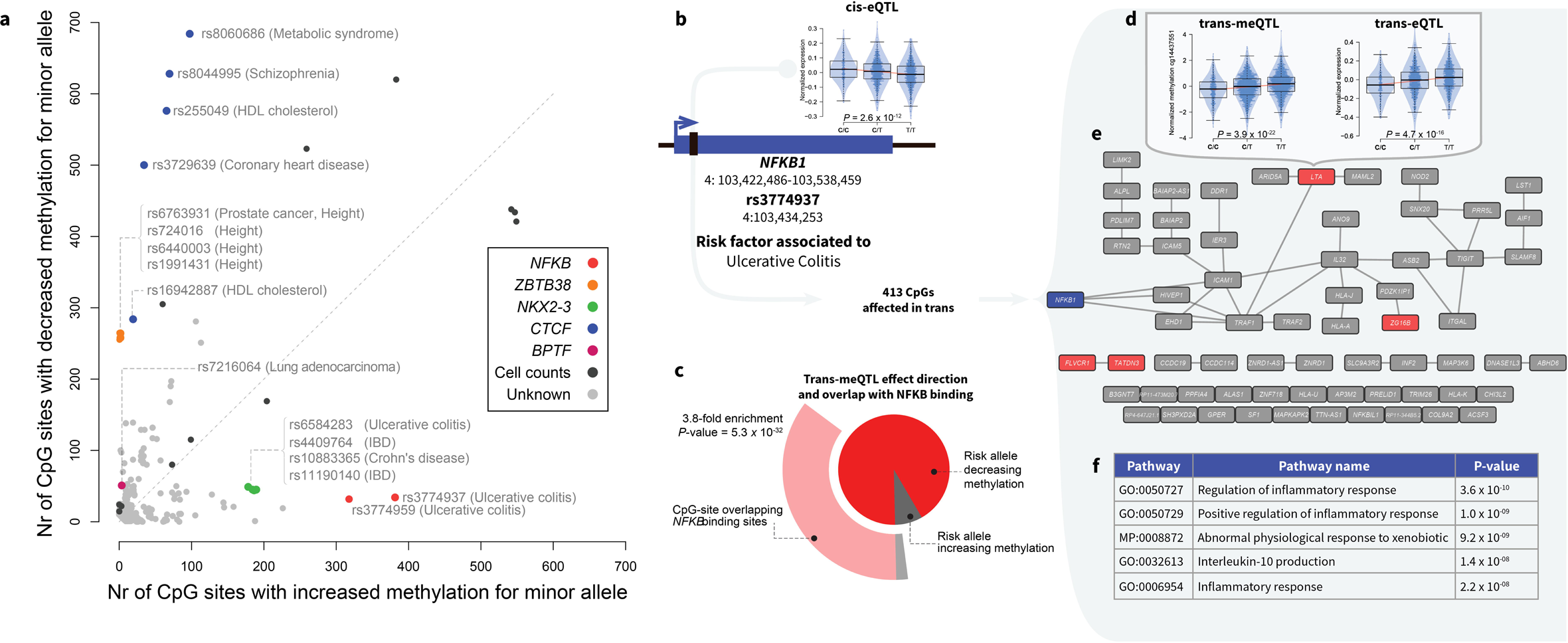
**a**, An imbalance in effect direction of *trans*-meQTLs implies involvement of transcription factors. Each dot represents a SNP with at least 10 *trans*-meQTL effects. The x-axis shows the number of *trans*-effects where the minor allele increases methylation, whereas the y-axis shows a decrease in methylation. SNPs with a multitude of effects of which many have the same allelic direction often exhibit evidence for a cis-eQTL on a transcription factor (colored dots), and an overrepresentation of CpGs *in trans* overlapping with binding sites for that transcription factor. **b**, Depiction of the *NFKB1* gene and rs3774937, associated with ulcerative colitis. The plot shows an increased expression of *NFKB1* for the risk allele C. **c**, In addition to influencing *NFKB1* expression, rs3774937 also influences DNA methylation at 413 CpGs *in trans*, decreasing methylation levels at 93% of affected CpG sites (dark grey). In addition, many of the CpG sites (37.3%) overlap with NFKB binding sites (3.8-fold enrichment, *P*-value = 5.3 *×* 10^−32^), shown in the outer chart. **d**, Illustrations of meQTL (left plot) and eQTL effects (right plot) of rs3774937 *in trans*. Only SNP-gene combinations were tested where the gene was associated with one of the 413 CpGs with a *trans*-meQTL. **e**, Gene network of the eQTM genes associated with 72 of the 413 CpGs (17.4%), that are showing a *trans*-meQTL (red). NFKB is depicted in blue. Genes also showing evidence for a *trans*-eQTL effect are shown in red. **f**, Top pathways as identified by enrichment method DEPICT for which many of the genes in **e** were overrepresented. Many of the identified pathways were inflammation-related, in line with the inflammatory nature of ulcerative colitis.

In order to explore this mechanism further, we combined ChIP-seq data on TF binding at CpGs and *cis*-expression effects of SNPs to directly examine the involvement of TFs in mediating *trans*-meQTLs. Among trait-associated SNPs influencing at least 10 CpGs *in trans* (n=305), we identified 13 *trans*-meQTL SNPs with strong support for a role of TFs (Figure 4a).

The most striking example was a locus on chromosome 4 (Figure 4b), where two SNPs (rs3774937 and rs3774959, in strong LD) were associated with ulcerative colitis (UC)^26^. Top SNP rs3774937 was associated with differential DNA methylation at 413 CpG sites across the genome, 92% of which showed the same direction of the effect, i.e. lower methylation associated with the risk allele (binomial P=2.72*×*10^−69^). Of those 380 CpG sites with lower methylation, 147 (38.7%) overlap with a nuclear factor kappaB (NFKB) transcription factor binding site (2.75-fold enrichment, *P* = 5.3*×*10^−32^), as based on ENCODE NFKB ChIP-seq data in blood cell types (Figure 4c). Three motif enrichment analyses (Homer, PWMEnrich, DEEPbind)^21–23^ also revealed an enrichment of NFKB binding motifs for the 413 CpG sites thus corroborating the ChIP-seq results. Notably, SNP rs3774937 is located in the first intron of *NFKB1* and we found that the risk allele was associated with higher *NFKB1* expression (Figure 4a). Of the 413 *trans*-CpGs, 64 were eQTMs and revealed a coherent gene network (Figure 4d) that was enriched for immunological processes related to *NFKB1* function^27^ (Figure 4e). Taken together, these results support the idea that the rs3774937 UC risk allele decreases DNA methylation *in trans* by increasing *NFKB1* expression *in cis*.

The same analysis approach indicated that the *trans*-methylation effects of rs8060686 (linked to various phenotypes including metabolic syndrome^28^ and coronary heart disease^29^, and affecting 779 *trans*-CpGs) were due to CTCF which mapped 315 kb from rs8060686. We observed a strong CTCF ChIP-seq enrichment with 603/779 trans-CpGs overlapping with CTCF binding (P =1.6×10^−232^) and enriched CTCF motifs (Figure 4a and Extended Data Fig. 5). Of these *trans*-CpGs, only13 have been observed previously in lymphocytes^18^. We observed that the risk allele increased DNA methylation *in trans* by decreasing *CTCF* gene expression *in cis*.

We found another example of this phenomenon: 228 *trans*-meQTL effects of 4 SNPs on chromosome 10, mapping near *NKX2–3* and implicated in inflammatory bowel disease^26^, were strongly enriched for NKX2 transcription factor motifs and associated with *NKX2–3* expression. The risk alleles decreased DNA methylation *in trans* at NKX2–3 binding sites by increasing *NKX2–3* gene expression *in cis* (Extended Data Fig. 6).

One height locus^30^ contained 4 SNPs which influence 267 *trans*-CpGs and implicate *ZBTB38* (Extended Data Fig. 7). In contrast to the aforementioned TFs that are transcriptional activators, *ZBTB38* is a transcriptional repressor^31,32^ and its expression was positively correlated with methylation *in trans*, in line with our observation that eQTMs in repressed regions are enriched for positive correlations. Finally, the *trans*-methylation effects of rs7216064 (64 *trans*-CpGs), associated with lung carcinoma^33^, preferentially occurred at regions binding CTCF, while the SNP was located in the *BPTF* gene, known to occupy CTCF binding sites^34^ (Extended Data Fig. 8).

The possibility to link *trans*-meQTL effects to an association of TF expression in *cis* and concomitant differential methylation in *trans* at the respective binding site is limited to TFs for which ChIP-seq data or motif information is available. In order to make inferences on TFs for which such data is not yet available, we ascertained whether *trans*-meQTLs SNPs were more often affecting TF gene expression in *cis* as compared with SNPs that were not giving *trans*-meQTLs. We observed that 13.1% of the GWAS SNPs that gave *trans*-meQTLs also affect TF gene expression in *cis*, whereas only 4.5% of the GWAS SNPs that do not give *trans*-meQTLs affect TF gene expression in cis (Fisher’s exact P = 6.6 × 10−13).

Here we report that one third of known disease- and trait-associated SNPs has downstream methylation effects *in trans*, often affecting multiple regions across the genome. The biological mechanism underlying *trans*-meQTLs often involves a local effect on the transcriptional activity of nearby TFs that affects DNA methylation at distal binding sites of the corresponding TFs. The direction of downstream methylation effects is remarkably consistent for each SNP and indicates that decreased DNA methylation is a signature of increased binding of transcriptional activators. Our study reveals previously unrecognized functional consequences of disease variants in non-coding regions. These can be looked up online (http://www.genenetwork.nl/biosqtlbrowser/), and provide leads for experimental follow-up.

## Methods

### Cohort descriptions

The five cohorts used in our study are described briefly below. The number of samples per cohort and references to full cohort descriptions can be found in Extended data table 1.

### CODAM

The Cohort on Diabetes and Atherosclerosis Maastricht^13^ (CODAM) consists of a selection of 547 subjects from a larger population-based cohort.^35^ Inclusion of subjects into CODAM was based on a moderately increased risk to develop cardiometabolic diseases, such as type 2 diabetes and/or cardiovascular disease. Subjects were included if they were of Caucasian descent and over 40 yrs of age and additionally met at least one of the following criteria: increased BMI (>25), a positive family history of type 2 diabetes, a history of gestational diabetes and/or glycosuria, or use of anti-hypertensive medication.

### LifeLines-DEEP

The LifeLines-DEEP (LLD) cohort^12^ is a sub-cohort of the LifeLines cohort.^36^ LifeLines is a multi-disciplinary prospective population-based cohort study examining the health and health-related behaviours of 167,729 individuals living in the northern parts of The Netherlands using a unique three-generation design. It employs a broad range of investigative procedures assessing the biomedical, socio-demographic, behavioural, physical and psychological factors contributing to health and disease in the general population, with a special focus on multi-morbidity and complex genetics. A subset of 1,500 LifeLines participants also take part in LLD^12^. For these participants, additional molecular data is generated, allowing for a more thorough investigation of the association between genetic and phenotypic variation.

### LLS

The aim of the Leiden Longevity Study^14^ (LLS) is to identify genetic factors influencing longevity and examine their interaction with the environment in order to develop interventions to increase health at older ages. To this end, long-lived siblings of European descent were recruited together with their offspring and their offspring’s partners, on the condition that at least two long-lived siblings were alive at the time of ascertainment. For men the age criteria was 89 or older, for women age 91 or over. These criteria led to the ascertainment of 944 long-lived siblings from 421 families, together with 1,671 of their offspring and 744 partners.

### NTR

The Netherlands Twin Register^15,37,38^ (NTR) was established in 1987 to study the extent to which genetic and environmental influences cause phenotypic differences between individuals. To this end, data from twins and their families (nearly 200,000 participants) from all over the Netherlands are collected, with a focus on health, lifestyle, personality, brain development, cognition, mental health, and aging. In NTR Biobank^15^ samples for DNA, RNA, cell lines and for biomarker projects have been collected.

### RS

The Rotterdam Study^16^ is a single-centre, prospective population-based cohort study conducted in Rotterdam, the Netherlands^16^. Subjects were included in different phases, with a total of 14,926 men and women aged 45 and over included as of late 2008. The main objective of the Rotterdam Study is to investigate the prevalence and incidence of and risk factors for chronic diseases to contribute to a better prevention and treatment of such diseases in the elderly.

## Genotype data

### Data generation

Genotype data was generated for each cohort individually. Details on the methods used can be found in the individual papers (CODAM: van Dam et al.^35^; LLD: Tigchelaar et al.^12^; LLS: Deelen et al.^39^, 2014; NTR: Willemsen et al.^15^; RS: Hofman et al.^16^).

### Imputation and QC

For each cohort separately, the genotype data were harmonized towards the Genome of the Netherlands^40^ (GoNL) using Genotype Hamonizer^41^ and subsequently imputed per cohort using Impute2^42^ using GoNL^43^ reference panel^43^ (v5). Quality control was also performed per cohort. We removed SNPs with an imputation info-score below 0.5, a HWE *P*-value smaller than 10^−4^, a call rate below 95% or a minor allele frequency smaller than 0.05. These imputation and filtering steps resulted in 5,206,562 SNPs that passed quality control in each of the datasets.

## Methylation data

### Data generation

For the generation of genome-wide DNA methylation data, 500 ng of genomic DNA was bisulfite modified using the EZ DNA Methylation kit (Zymo Research, Irvine, California, USA) and hybridized on Illumina 450k arrays according to the manufacturer’s protocols. The original IDAT files were generated by the Illumina iScan BeadChip scanner. We collected methylation data for a total of 3,841 samples. Data was generated by the Human Genotyping facility (HugeF) of ErasmusMC, the Netherlands (www.glimDNA.org).

### Probe remapping and selection

We remapped the 450K probes to the human genome reference (HG19) to correct for inaccurate mappings of probes and identify probes that mapped to multiple locations on the genome. Details on this procedure can be found in Bonder et al. (2014)^44^. Next, we removed probes with a known SNP (GoNL, MAF > 0.01) at the single base extension (SBE) site or CpG site. Lastly, we removed all probes on the sex chromosomes, leaving 405,709 high quality methylation probes for the analyses.

### Normalization and QC

Methylation data was directly processed from IDAT files resulting from the Illumina 450k array analysis, using a custom pipeline based on the pipeline developed by Tost & Toulemat^45^. First, we used methylumi^46^ to extract the data from the raw IDAT files. Next, we performed quality control checks on the probes and samples, starting by removing the incorrectly mapped probes. We checked for outlying samples using the first two principal components (PCs) obtained using principal component analysis (PCA). None of the samples failed our quality control checks, indicating high quality data. Following quality control, we performed background correction and probe type normalization as implemented in DASEN^47^. Normalization was performed per cohort, followed by quantile normalization on the combined data to normalize the differences per cohort. The next step in quality control consisted of identifying potential sample mix-ups between genotype and DNA methylation data. Using mix-up mapper^48^, we detected and corrected 193 mix-ups. Lastly, in order to correct for known and unknown confounding sources of variation in the methylation data and to give us more power to detect meQTLs, we removed the first components which were not affected by genetic information, the 22 first PCs, from the methylation data using methodology we have successfully used in *trans*-eQTL^3^’^49^ and meQTL analyses before^44^.

### RNA sequencing

Total RNA from whole blood was deprived of globin using Ambion’s GLOBIN clear kit and subsequently processed for sequencing using Illumina’s Truseq version 2 library preparation kit. Paired-end sequencing of 2×50bp was performed using Illumina’s Hiseq2000, pooling 10 samples per lane. Finally, read sets per sample were generated using CASAVA, retaining only reads passing Illumina’s Chastity Filter for further processing. Data was generated by the Human Genotyping facility (HugeF) of ErasmusMC, the Netherlands (www.glimDNA.org).

Initial QC was performed using FastQC^50^ (v0.10.1), removal of adaptors was performed using cutadapt^51^ (v1.1), and Sickle^52^ (V1.2) [2] was used to trim low quality ends of the reads (min length 25, min quality 20). The sequencing reads were mapped to human genome (HG19) using STAR^53^ v2.3.125. Gene expression quantification was performed by HTseq-count. The gene definitions used for quantification were based on Ensmble version 71, with the extension that regions with overlapping exons were treated as separate genes and reads mapping within these overlapping parts did not count towards expression of the normal genes.

Expression data on the gene level were first normalized using Trimmed Mean of M-values^54^. Then expression values were log2 transformed, gene and sample means were centred to zero. To correct for batch effects, PCA was run on the sample correlation matrix and the first 25 PCs were removed using methodology that we have use for eQTL analyses before^49,55^. More details are provided in Zhernakova et al (in preperation).

### *Cis*-meQTL mapping

In order to determine the effect of nearby genetic variation on methylation levels (*cis*-meQTLs), we performed *cis*-meQTL mapping using 3,841 samples for which both genotype data and methylation data were available. To this end, we calculated the Spearman rank correlation and corresponding *P*-value for each CpG-SNP pair in each cohort separately. We only considered CpG-SNP pairs located no further than 250kb apart. The *P*-values were subsequently transformed into a Z-score for meta-analysis. To maximize the power of meQTL detection, we performed a meta-analysis over all datasets by calculating an overall, joint *P*-value using a weighted Z-method. A comprehensive overview of this method has been described previously^55^. To detect all possible independent SNPs regulating methylation at a single CpG-site we regressed out all primary *cis*-meQTL effects and then ran *cis*-meQTL mapping for the same CpG-site to find secondary *cis*-meQTL. We repeated that in a stepwise fashion until no more independent *cis*-meQTL were found.

To filter out potential false positive *cis*-meQTLs caused by SNPs affecting the binding of a probe on the array, we filtered the *cis*-meQTLs effects by removing any CpG-SNP pair for which the SNP was located in the probe. In addition, all other CpG-SNP pairs for which the SNP was outside the probe, but in LD (R^2^ > 0.2 or D’ > 0.2) with a SNP inside the probe were also removed. We tested for LD between SNPs in the probe and in the surrounding *cis* area in the individual genotype datasets, as well as in GoNL v5, in order to be as strict as possible in marking a QTL as true positive.

To correct for multiple testing, we empirically controlled the false discovery rate (FDR) at 5%. For this, we compared the distribution of observed *P*-values to the distribution obtained from performing the analysis on permuted data. Permutation was done by shuffling the sample identifiers of one data set, breaking the link between, e.g., the genotype data and the methylation or expression data. We repeated this procedure 10 times to obtain a stable distribution of *P*-values under the null distribution. The FDR was determined by only selecting the strongest effect per CpG^55^ in both the real analysis and in the permutations (i.e. probe level FDR < 5%).

### *Cis*-eQTL mapping

For a set of 2,116 BIOS samples we had also generated RNA-seq data. We used this data to identify *cis*-eQTLs. *Cis*-eQTL mapping was performed using the same method as *cis*-meQTL mapping. Details on these eQTLs will be described in a separate paper (Zhernakova *et al*, in preparation).

## Expression quantitative trait methylation (eQTM) analysis

To identify associations between methylation levels and expression levels of nearby genes (*cis*-eQTMs), we first corrected our expression and methylation data for batch effects and covariates by regressing out the PCs and regressing out the identified *cis*-meQTLs and *cis*-eQTLs, to ensure only relationships between CpG sites and gene expression levels would be detected that were not attributable to particular genetic variation or batch effects. We mapped eQTMs in a window of 250Kb around the TSS of a transcript. Further statistical analysis was identical to the *cis*-meQTL mapping. For this analysis we were able to use a total of 2,101 samples for which both genetic, methylation and gene expression data was available. To correct for multiple testing we controlled the FDR at 5%, the FDR was determined by only selecting the strongest effect per CpG^55^ in both the real analysis and in the permutations.

### *Trans*-meQTL mapping

To identify the effects of distal genetic variation with methylation (*trans*-meQTLs) we used the same 3,841 samples that we had used for *cis*-meQTL mapping. To focus our analysis and limit the multiple testing burden, we restricted our analysis to SNPs that have been previously found to be significantly correlated to traits and diseases at a *P* < 5×10^−8^. We extracted these SNPs from the NHGRI genome-wide association study (GWAS) catalogue, used recent GWAS studies not yet in the NHGRI GWAS catalogue and studies on the Immunochip and Metabochip platform that are not included in the NHGRI GWAS catalogue (Extended Data Table 1). We compiled this list of SNPs in December 2014. Per SNP we only investigated CpG sites that mapped at least 5 Mb from the SNP or on other chromosomes. Before mapping *trans*-meQTLs, we regressed out the identified *cis*-meQTLs to increase the statistical power of *trans*-meQTL detection (as done previously for *trans*-eQTLs^3^) and to avoid designating an association as *trans* that may be due to long-range LD (e.g. within the HLA region). To ascertain the stability of the *trans*-meQTLs we also performed the *trans*-mapping on the non-corrected data and the methylation data corrected for cell-type proportions. In addition, we performed meQTL mapping on SNPs known to influence the cell type proportions in blood^19,20^.

To filter out potential false positive *trans*-meQTLs due to cross-hybridization of the probe, we remapped the methylation probes with very relaxed settings, identical to Westra et al.^55^, with the difference that we only accepted mappings if the last bases of the probe including the SBE site were mapped accurately to the alternative location. If the probe mapped within our minimal *trans*-window, 5 Mb from the SNP, we removed the effect as being a false positive *trans*-meQTL.

We controlled for multiple testing by using 10 permutations. We controlled the false-discovery rate at 5%, identical to the aforementioned *cis*-meQTL analysis.

### *Trans*-eQTL mapping

To check if the *trans*-meQTL effects can also be found back on gene expression levels, we annotated the CpGs with a *trans*-meQTL to genes using our eQTMs. Using the 2,101 samples for which both genotype and gene expression data were available, we performed *trans*-eQTL mapping, associating the SNPs known to be associated with DNA methylation in *trans* with their corresponding eQTM genes.

## Annotations and enrichment tests

Annotation of the CpGs was performed using Ensembl^56^ (v70), UCSC Genome Browser^57^ and data from the Epigenomics Roadmap Project.^58^ We used the Epigenomics Roadmap annotation for the SBE site of the methylation site for all 27 blood cell types. We chose to use both the histone mark information and the chromatin marks in blood-related cell types only, as generated by the Epigenomics Roadmap Project. Summarizing the information over the 27 blood cell types was done by counting presence of histone-marks in all the cell types and scaling the abundance, i.e. if the mark is bound in all cell types the score would be 1 if it would be present in none of the blood cell types the score would be 0.

To calculate enrichment of meQTLs or eQTMs for any particular genomic context, we used logistic regression because this allows us to account for covariates such as CpG methylation variation. For *cis*-meQTLs, we used the variability of DNA methylation, the number of SNPs tested, and the distance to the nearest SNP per CpG as covariates. For all other analyses we used only the variability in DNA methylation as a covariate.

Next to annotation data from the Epigenomics Roadmap project, we used transcription factor ChIP-seq data from the ENCODE-project for blood-related cell lines. For every CpG site, we determined if there was an overlap with a ChIP-seq signal and performed a Fisher exact test to determine whether the *trans*-meQTL probes associated with the SNP in the transcription factor region of interest were more often overlapping with a ChIP-seq region than the other *trans*-meQTL probes. We collected all transcription factor called narrow peak files from the UCSC genome browser to perform the enrichments.

Enrichment of known sequence motifs among *trans*-CpGs was assessed by PWMEnrich^22^ package in R, Homer^59^ and DEEPbind^23^. For PWMEnrich hundred base pair sequences around the interrogated CpG site were used, and as a background set we used the top CpGs from the 50 permutations used to determine the FDR threshold of the *trans*-meQTLs. For Homer the default settings for motif enrichment identification were used, and the same CpGs derived from the permutations were used as a background. For DEEPbind we used both the permutation background like described for Homer and the permutations background as described for PWMEnrich.

Using data published by Rao *et al*.^10^ we were able to intersect the *trans*-meQTLs with information about the 3D structure of the human genome. For the annotation, we used the combined Hi-C data for both inter- and intra-chromosomal data at 1Kb and the quality threshold of E30 in the GM12878 lymphoblastoid cell line. Both the *trans*-meQTL SNP and *trans*-meQTL probes were put in the relevant 1Kb block, and for these blocks we looked up the chromosomal contact value in the measurements by Rao et al. Surrounding the *trans*-meQTLs SNPs, we used a LD window that spans maximally 250Kb from the *trans*-meQTL SNP and had a minimal R^2^ of 0. 8. If a Hi-C contact between the SNP block and the CpG-site was indicated, we flagged the region as a positive for Hi-C contacts. As a background, we used the combinations found in our 50 permutated *trans*-meQTL analyses, taking for each permutation the top trans-meQTLs that were similar in size to the real analysis. This permitted us to empirically determine whether there were significantly more Hi-C interactions in the real data as compared to the permutations.

### eQTM direction prediction

We predicted the direction of the eQTM effects using both a decision tree and a naïve Bayes model (as implemented by Rapid-miner^60^ v6.3). We built the models on the strongest eQTMs (i.e. those identified at a very stringent FDR <9.73×10^−6^). For the decision tree we used a standard cross-validation set-up using 20 folds. For the naive Bayes model we used a double loop cross-validation: performance was evaluated in the outer loop using 20-fold cross-validation, while feature selection (using both backward elimination and forward selection) took place in the inner loop using 10-fold cross-validation. Details about the double-loop cross-validation can be found in Ronde et al.^61^. During the training of the model, we balanced the two classes making sure we had an equal number of positively correlating and negatively correlating CpG-gene combinations, by randomly sampling a subset of the overrepresented negatively correlating CpG-gene combination group. We chose to do so to circumvent labelling al eQTMs as negative, since this is the class were the majority of the eQTMs are in.

In the models we used annotation from the CpG-site, namely: overlap with epigenomics roadmap chromatin states, histone marks and relations between the histone marks, GC content surrounding the CpG-site and relative locations from the CpG-site to the transcript.

## DEPICT

To investigate whether there was biological coherence in the *trans*-meQTLs identified, we performed gene-set enrichment analysis for each genetic risk factor that was showing at least 10 *trans*-meQTL effects. To do so, we adapted DEPICT^27^, a pathway enrichment analysis method that we previously developed for GWAS. Instead of defining loci with genes by using top associated SNPs, we used the eQTM information to link CpGs to genes. Within DEPICT gene set enrichment, significance is determined by using matched sets of permuted loci (in terms of numbers of genes per locus) that have been identified using simulated GWAS. Subsequent pathway enrichment analysis was conducted as described before, and significance was determined by controlling the false discovery rate at 5%.

## Acknowledgements

This work was performed within the framework of the Biobank-Based Integrative Omics Studies (BIOS) Consortium funded by BBMRI-NL, a research infrastructure financed by the Dutch government (NWO 184.021.007). Samples were contributed by LifeLines (http://lifelines.nl/lifelines-research/general), the Leiden Longevity Study (http://www.healthy-ageing.nl; http://www.leidenlangleven.nl), the Netherlands Twin Registry (NTR: http://www.tweelingenregister.org), the Rotterdam studies, (http://www.erasmus-epidemiology.nl/rotterdamstudy), the Genetic Research in Isolated Populations program (http://www.epib.nl/research/geneticepi/research.html#gip), the Codam study (http://www.carimmaastricht.nl/) and the PAN study (http://www.alsonderzoek.nl/). We thank the participants of all aforementioned biobanks and acknowledge the contributions of the investigators to this study (Supplemental Acknowledgements). This work was carried out on the Dutch national e-infrastructure with the support of SURF Cooperative.

## Author contributions

BTH, PACtH, JBJvM, AI, RJ and LF formed the management team of the BIOS consortium. DIB, RP, JVD, JJH, MMJVG, CDAS, CJHvdK, CGS, CW, LF, AZ, EFG, PES, MB, JD, DvH, JHV, LHvdB, CMvD, BAH, AI, AGU managed and organized the biobanks. JBJvM, PMJ, MV, HEDS, MV, RvdB, JvR and NL generated RNA-seq and Illumina 450k data. HM, MvI, MvG, JB, DVZ, RJ, PvtH, PD, IN, PACtH, BTH and MM were responsible for data management and the computational infrastructure. MJB, RL, MV, DVZ, RS, IJ, MvI, PD, FvD, MvG, WA, SMK, MAS, EWvZ, RJ, PACtH, LF and BTH performed the data analysis. MJB, RL, LF and BTH drafted the manuscript.

## Data availability

All results can be queried using our dedicated QTL browser: http://genenetwork.nl/biosqtlbrowser/. Raw data was submitted to the European Genome-phenome Archive (EGA)

